# SIGNAL-seq: Multimodal Single-cell Inter- and Intra-cellular Signalling Analysis

**DOI:** 10.1101/2024.02.23.581433

**Authors:** James W. Opzoomer, Rhianna O’Sullivan, Jahangir Sufi, Ralitsa Madsen, Xiao Qin, Ewa Basiarz, Christopher J. Tape

## Abstract

We present SIGNAL-seq (Split-pool Indexing siG-Nalling AnaLysis by sequencing): a multiplexed split-pool combinatorial barcoding method that simultaneously measures RNA and post-translational modifications (PTMs) in fixed single cells from 3D models. SIGNAL-seq PTM measurements are equivalent to mass cytometry and RNA gene detection is analogous to split-pool barcoding scRNA-seq. By measuring both mRNA ligand-receptor pairs and PTMs in single cells, SIGNAL-seq can simultaneously uncover inter- and intra-cellular regulation of tumour microenvironment plasticity.

## Introduction

Cells process extra-cellular signals via intra-cellular protein post-translational modifications (PTMs) to regulate gene expression [1]. Common PTMs include protein phosphorylation, acetylation, methylation, and protease cleavage — each regulating a range of cellular phenotypes.

PTM signalling can be measured in single cells via anti-PTM antibodies using flow or mass cytometry, but these methods are limited to <40 PTMs per cell due to finite fluorophores or monoisotopic rare-earth metals [2]. Single-cell antigen detection can be expanded indefinitely using DNA-oligonucleotide conjugated antibodies that encode protein abundance as sequenceable antibody-derived tags (ADTs). ADTs are typically used to analyse extra-cellular proteins alongside RNA on droplet-based microfluidic single-cell RNA sequencing (scRNA-seq) platforms (e.g. CITE-seq [3]). ADTs can also be used to detect transcription factors via singlenucleus RNA sequencing (snRNA-seq) [4] and scRNAseq [5], phospho-proteins and mRNA at low-throughput with reversible fixation [6], and phospho-proteins at the expense of poor mRNA coverage [7, 8]. However, despite the fundamental link between PTM signalling and transcriptional responses, no method can simultaneously measure both intra-cellular PTM signalling and broad polyadenylated and non-polyadenylated RNA profiles in a scalable, instrument-independent, and cost-effective manner (Table 1).

**Table 1.** Comparison of SIGNAL-seq to Published Multimodal Single-cell Technologies. * = Published average RNA genes per cell: + <100; ++ <1,000; +++ >2,000. ** = Cell throughput per experiment based on foundational technology: + <1,000 cells; ++ 1,000-10,000 cells; +++ 10,000->100,000 cells. *** = Cost per 1M cells based on foundational technology (from Sziraki *et al*., 2023 [52]).

| Method | Foundation | Modalities | Genes/Cell* | Model Systems | Throughput** | Cost*** (\$/M) |
| --- | --- | --- | --- | --- | --- | --- |
| inCITE-seq [4] | Droplet (10X Genomics) | RNA<br>TF Proteins | +++ | 2D Cell Line<br>Tissue | ++ | 330K |
| NEAT-seq [5] | Droplet (10X Genomics) | RNA<br>TF Proteins<br>Chromatin Accessibility | +++ | 2D Cell Lines<br>PBMCs | ++ | 330K |
| RAID [6] | CELseq2 (Custom) | RNA (reversible fixation)<br>PTM (Phospho) | +++ | 2D Cell Lines | + | N/A |
| QuRIE-seq [7] | Droplet (Custom) | RNA<br>PTM (Phospho) | +<br>++ | B-cells<br>Cell Lines | ++ | N/A |
| Phospho-seq [8] | Droplet (10X Genomics) | Chromatin Accessibility<br>PTM (Phospho)<br>Intra-cellular Proteins | None (requires bridging) | 2D Cell lines<br>3D Organoids | ++ | 330K |
| SPARC [53] | SMART-seq2 + PEA | RNA<br>Intra-cellular Proteins | +++ | 2D Cell Lines | + | N/A |
| DOGMA-seq [54] | Droplet (10X Genomics) | RNA<br>Chromatin Accessibility<br>Extra-cellular Protein | +++ | PBMCs | ++ | 330K |
| TEA-seq [55] | Droplet (10X Genomics) | RNA<br>Chromatin Accessibility<br>Extra-cellular Proteins | +++ | PBMCs | ++ | 330K |
| **SIGNAL-seq** | **Split-pool Barcoding** | **RNA (PolyA+rHex capture)**<br>**PTM (Phospho, Protease)**<br>**Intra-cellular Protein** | **+++** | **3D Organoids**<br>**3D Spheroids** | **+++** | **10K** |

Here, we describe SIGNAL-seq (Split-pool Indexing siGNalling AnaLysis by sequencing), a multimodal method that combines intra-cellular mass cytometry staining protocols with anti-PTM/protein ADTs and split-pool combinatorial barcoding to simultaneously measure intra-cellular proteins, cytoplasmic and nuclear PTMs, polyadenylated RNA, and non-polyadenylated RNA in tens of thousands of fixed single cells from solid-phase 3D cancer models.

We demonstrate that SIGNAL-seq PTM measurements are comparable to gold standard mass cytometry assays and transcriptomic gene detection is equivalent to Split Pool Ligation-based Transcriptome sequencing (SPLiT-seq) [9]. By simultaneously measuring mRNA ligand-receptor pairs, intra-cellular PTMs, and transcriptional responses in tens of thousands of single cells, SIGNAL-seq can assess both inter- and intra-cellular regulation of cell plasticity in a single assay. SIGNAL-seq does not require any custom instrumentation and can be established in any standard laboratory.

## Results

### SIGNAL-seq

SIGNAL-seq combines optimised intra-cellular staining workflows from mass cytometry [10, 11] with anti-PTM/protein ADTs and combinatorial split-pool barcoding scRNA-seq [9] to simultaneously measure PTMs, intra-cellular proteins, and RNA in fixed cells.

In SIGNAL-seq, we first fix single cells with formaldehyde to preserve PTMs and cell-states, and then permeabilise with Triton X-100 (full details in Methods). This sample preparation workflow is similar to standard intra-cellular flow and mass cytometry assays [10, 11]. To minimise non-specific background staining, permeabilised cells are blocked with BSA, dextran sulphate [4], and sheared salmon sperm DNA [5], and then stained with poly-A barcoded oligonucleotide-tagged antibodies targeting cytoplasmic and nuclear PTMs and intra- and extra-cellular proteins (previously optimised using mass cytometry [10, 11, 12, 13]).

Stained cells then enter a SPLiT-seq combinatorial barcoding workflow [9]. Unlike droplet-based scRNA-seq methods that are primarily designed to analyse live cells, combinatorial barcoding technologies such as SPLiT-seq [9] and sci-RNA-seq [14] are optimised to analyse RNA from fixed and permeabilised cells. SIGNAL-seq uses split-pool combinatorial barcoding to generate parallel NGS libraries for RNA and ADT-derived PTM/protein modalities from fixed and permeabilised cells. Through the use of both oligo-dT and random hexamer reverse transcription primers, SIGNAL-seq can measure both polyadenylated (poly-A) RNA and non-polyadenylated RNA (e.g. many short non-coding, long non-coding, and non-polyadenylated protein-coding transcripts). RNA and PTM/protein reads are then assigned single-cell identities via their split-pool cell barcodes and computationally integrated into a multimodal data matrix (Figure 1a).

**Figure 1.**
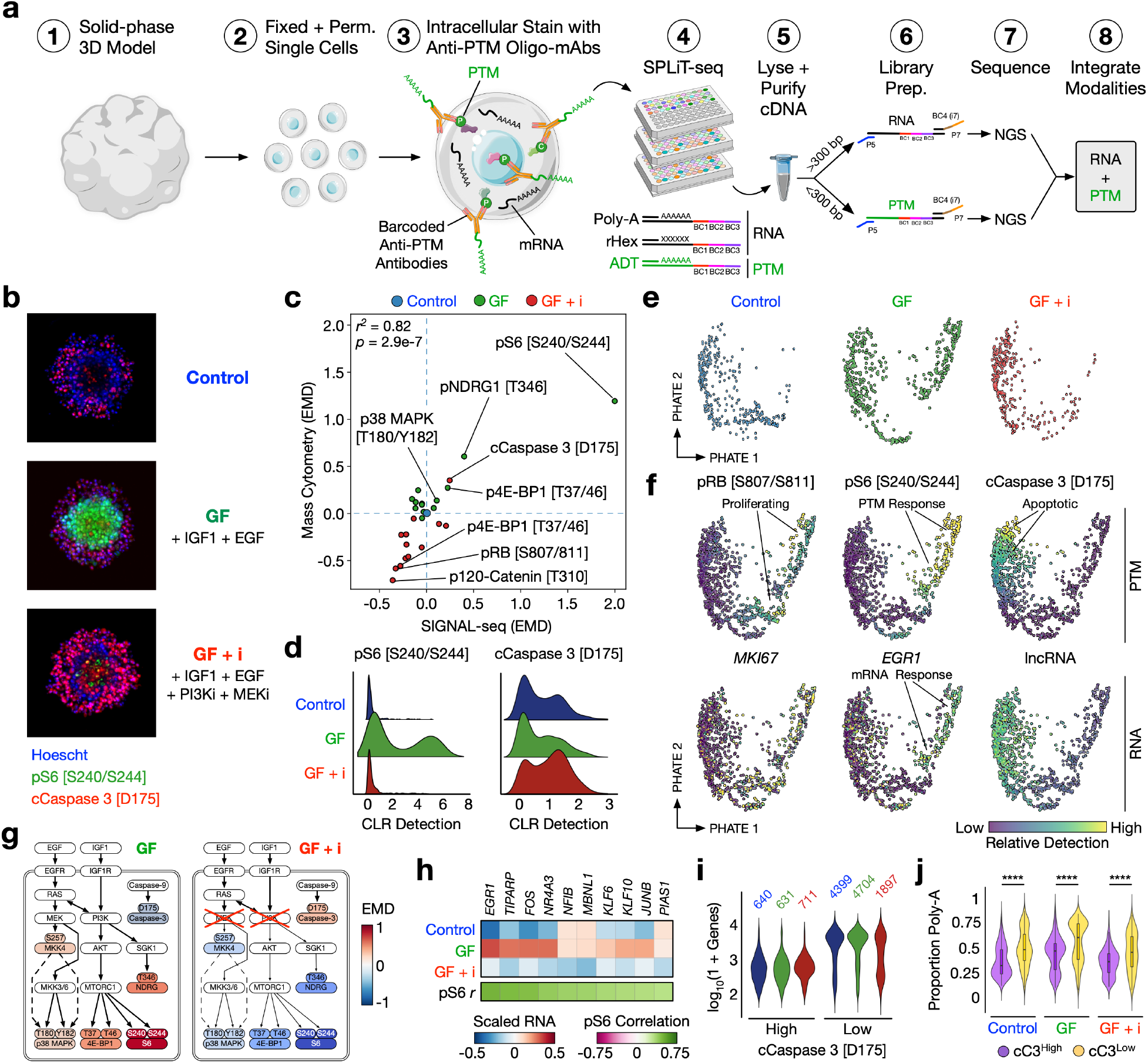
SIGNAL-seq, an intra-cellular staining protocol to detect proteins, PTM signalling and transcriptome in single-cells by combinatorial barcoding. **a)** SIGNAL-seq workflow. p, phosphorylation. c, cleavage. rHex, random hexamer. **b)** Cervical cancer (HeLa) 3D spheroids treated with either EGF and IGF1 growth factors (GF) or GF + inhibitors (GF + i), stained with pS6 [S240/S244] (green), cCaspase3 (cC3) [D175] (red) or Hoescht. Spheroids have a GF-responsive core (pS6^+^), and an apoptotic periphery (cC3^+^). **c)** Spheroids from b) analysed using SIGNAL-seq and mass cytometry demonstrate similar response to GF and GF + i. Earth mover’s distance (EMD) was calculated for each PTM relative to control spheroids. **d)** CLR (Centered log ratio transformation) detection of pS6 and cCaspase3 across each condition by SIGNAL-seq. **e)** Single-cell PHATE embeddings built with all cells from all conditions on the ADT modality, split by treatment. **f)** Single-cell PHATE embeddings of all conditions annotated by either PTM or RNA features. **g)** SIGNAL-seq EMD signalling model of GF and GF+i spheroids. **h)** Detection of top early EGF target gene responses by SIGNAL-seq and correlation to pS6 [S240/S244] activity. **i)** RNA gene counts per condition in cC3^High^ or cC3^Low^ cells. **j)** Proportion of poly-A (oligo-dT RT primed) transcripts per cell across conditions in cC3^High^ or cC3^Low^ populations. *t* -test = **** <0.0001.

To benchmark SIGNAL-seq in a defined 3D culture system, we stimulated HeLa spheroids for 30 minutes with IGF1 and EGF growth factors (GFs), +/-inhibitors of downstream kinases MEK (Trametinib) and class I PI3K (Pictilisib) (GFs + i). These conditions generate a dynamic range of signalling states [15], with spheroids containing a pS6 [S240/S244]^+^ IGF1 and EGF responsive core, and an cCaspase 3 [D175]^+^ apoptotic periphery when MEK and PI3K are inhibited (Figure 1b).

Following treatments, spheroids were fixed *in situ*, dissociated into single cells, and analysed in parallel by both thiol-reactive organoid barcoding *in situ* (TOB*is*) mass cytometry [10, 11] (Table 2) and SIGNAL-seq (Table 3). Earth mover’s distance (EMD) [16] analysis revealed that SIGNAL-seq and mass cytometry detect near identical PTM regulation following growth factor and inhibitor treatments (*r* ^2^ = 0.82) (Figure 1c).

**Table 2.** TOB*is* mass cytometry antibody panel used in TOB*is* HeLa spheroid experiments.

| Antigen | Metal | Clone | Supplier |
| --- | --- | --- | --- |
| pHistone H3 [S28] | <sup>89</sup>Y | HTA28 | BioLegend |
| Vimentin | <sup>111</sup>Cd | RV202 | BD Biosciences |
| CD326 (EpCAM) | <sup>113</sup>In | 9C4 | BioLegend |
| CK18 | <sup>114</sup>Cd | C-04 | Abcam |
| Pan-CK | <sup>115</sup>In | AE1/AE3 | Biolegend UK |
| GFP | <sup>116</sup>Cd | 5F12.4 | eBiosciences |
| pPDK1 [S241] | <sup>141</sup>Pr | J66-653.44.22 | BD Biosciences |
| cCaspase 3 [D175] | <sup>142</sup>Nd | D3E9 | CST |
| pMKK4 SEK1 [S257] | <sup>146</sup>Nd | C36C11 | CST |
| pBTK [Y511] | <sup>147</sup>Sm | 24a/BTK | BD Biosciences |
| p4E-BP1 [T37/46] | <sup>149</sup>Sm | 236B4 | CST |
| pRB [S807/811] | <sup>150</sup>Nd | J1112-906 | BD Biosciences |
| pNDRG1 [T346] | <sup>151</sup>Eu | D98G11 | CST |
| pAKT [T308] | <sup>152</sup>Sm | J1-223.371 | BD Biosciences |
| pNK-κB p65 [S529] | <sup>156</sup>Gd | K10-895.12.50 | BD Biosciences |
| pMKK3/pMKK6 [S189/207] | <sup>157</sup>Gd | D8E9 | CST |
| pP38 [T180/Y182] | <sup>158</sup>Gd | D3F9 | CST |
| pHistone H2A.X [S139] | <sup>162</sup>Dy | D7T2V | CST |
| p120 Catenin [T310] | <sup>164</sup>Dy | 22/p120 (pT310) | BD Biosciences |
| pS6 [S240/244] | <sup>172</sup>Yb | D68F8 | CST |
| CD44 | <sup>175</sup>Lu | IM7 | BioLegend |
| Cyclin B1 | <sup>176</sup>Yb | GNS-11 | BD Biosciences |
| meHistone H3 [K4] | <sup>209</sup>Bi | C64G9 | CST |

**Table 3.**
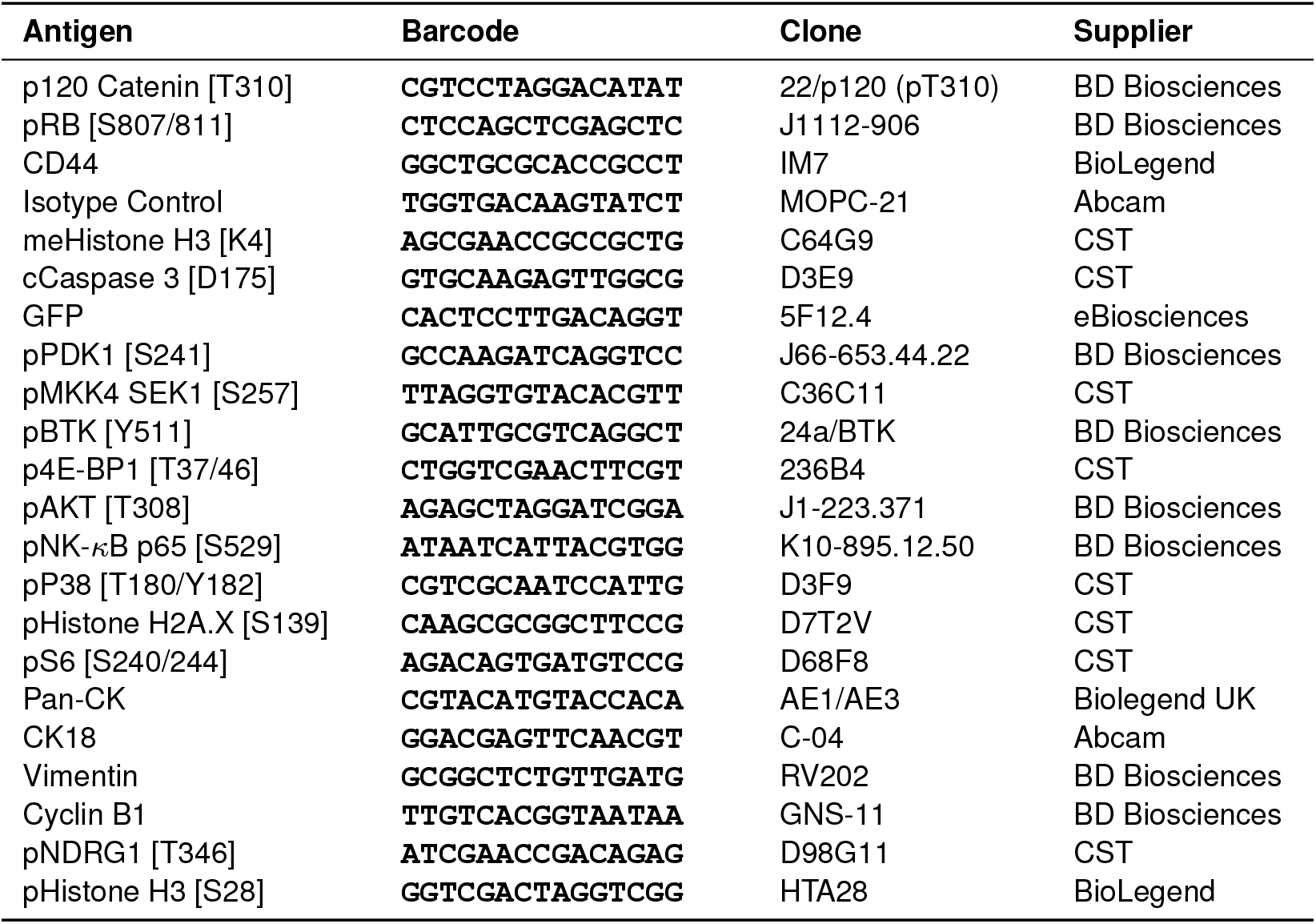
SIGNAL-seq antibody panel used in HeLa spheroid experiments.

SIGNAL-seq accurately identified a subpopulation pS6 [S240/S244] response to EGF and IGF1 and increased cCaspase 3 [D175] following MEK and PI3K inhibition (Figure 1d) (Figure S1a). Crucially, SIGNAL-seq detected intra-cellular regulation of growth factor signalling via both upstream PTMs (e.g. pS6 [S240/S244], pNDRG1 [T346], p4E-BP1 [T37/T46]) and downstream mRNA (e.g. *EGR1, FOS, JUNB*). IGF1 and EGF responsive cells were generally in the cell-cycle — again identified by both PTMs (pRB [S807/S811]^+^) and mRNA (*MKI67* ^+^) (Figure 1e-g). By contrast, inhibitor treated cells were largely apoptotic (cCaspase 3 [D175]^+^) (Figure S1b) and have high levels of long non-coding RNA (lncRNA). Despite the short 30 minute growth factor stimulation, SIGNAL-seq detected transcriptional up-regulation of several canonical growth factor early-response target genes [17] (Figure 1h) (Figure S1c-f). These results demonstrate that SIGNAL-seq can analyse how extra-cellular ligands regulate intra-cellular PTM signals to control RNA gene expression in 3D models.

Parallel analysis of PTMs and RNA enabled a direct comparison of cellular features captured by each modality. For example, we found that RNA detection was substantially higher in viable cells (cCaspase 3 [D175]^Low^) compared to apoptotic cells (cCaspase3 [D175]^High^) (Figure 1i) (Figure S1b). SIGNAL-seq identified 4,399 median genes / cell in viable serum-starved control spheroids — comparable to 3,500-5,500 median genes / cell from SPLiT-seq analysis of 2D cells grown in serum [9] (Figure S1g, h). SIGNAL-seq can also measure a broad array of both processed and unprocessed transcripts via poly-A and random hexamer capture primers. We found that cCaspase3 [D175]^High^ cells contain a lower relative proportion of poly-A mRNA in their transcriptome compared to cCaspase 3 [D175]^Low^ cells across all conditions (Figure 1j). This is in agreement with observations that apoptotic cells undergo global poly-A mRNA targeted degradation during apoptosis [18]. Comparison of pRB [S807/S811] against cell-cycle gene expression signatures [19] also demonstrated that RNA can be used to approximate cell-cycle phase (Figure S1i).

In summary, by combining mass cytometry staining protocols with anti-PTM ADTs and split-pool combinatorial barcoding, SIGNAL-seq can measure extra-cellular regulation of intra-cellular PTM signalling and downstream RNA expression. SIGNAL-seq PTM measurements are equivalent to mass cytometry and RNA detection is analogous to SPLiT-seq.

### SIGNAL-seq Analysis of Tumour Microenvironment Drug Response Plasticity

In solid-tumours, both inter-cellular signalling from the tumour microenvironment (TME) and anti-cancer drug treatments can regulate intra-cellular PTM signalling networks in cancer cells. Deregulated PTM signalling can then alter cancer cell gene expression to drive phenotypic plasticity [20]. For example, in colorectal cancer (CRC) inter-cellular signalling from cancer associated fibroblasts (CAFs) can regulate cancer cell PTM signalling to polarise cancer cells from a chemosensitive proliferative colonic stem cell (proCSC) fate to a chemorefractory revival stem cell (revCSC) fate [12, 21].

By measuring RNA and PTMs in fixed single cells, we hypothesised that SIGNAL-seq could perform inter- and intra-cellular signalling analysis of TME-driven drug response plasticity in a single assay. To test this, we cultured CRC patient derived organoids (PDOs) +/-CAFs in 3D, and treated +/-SN-38 chemotherapy (Figure 2a). This patient-derived 3D model system provides a dynamic range of signalling and cell-fates driven by both inter-cellular communication and chemotherapy. Fixed PDO-CAF single cells were analysed by SIGNAL-seq using an expanded ADT panel (Table 4) that included additional oligo-tagged antibodies against celltype-specific intra-cellular proteins and the DNA-damage marker pHH2AX [S139].

**Table 4.**
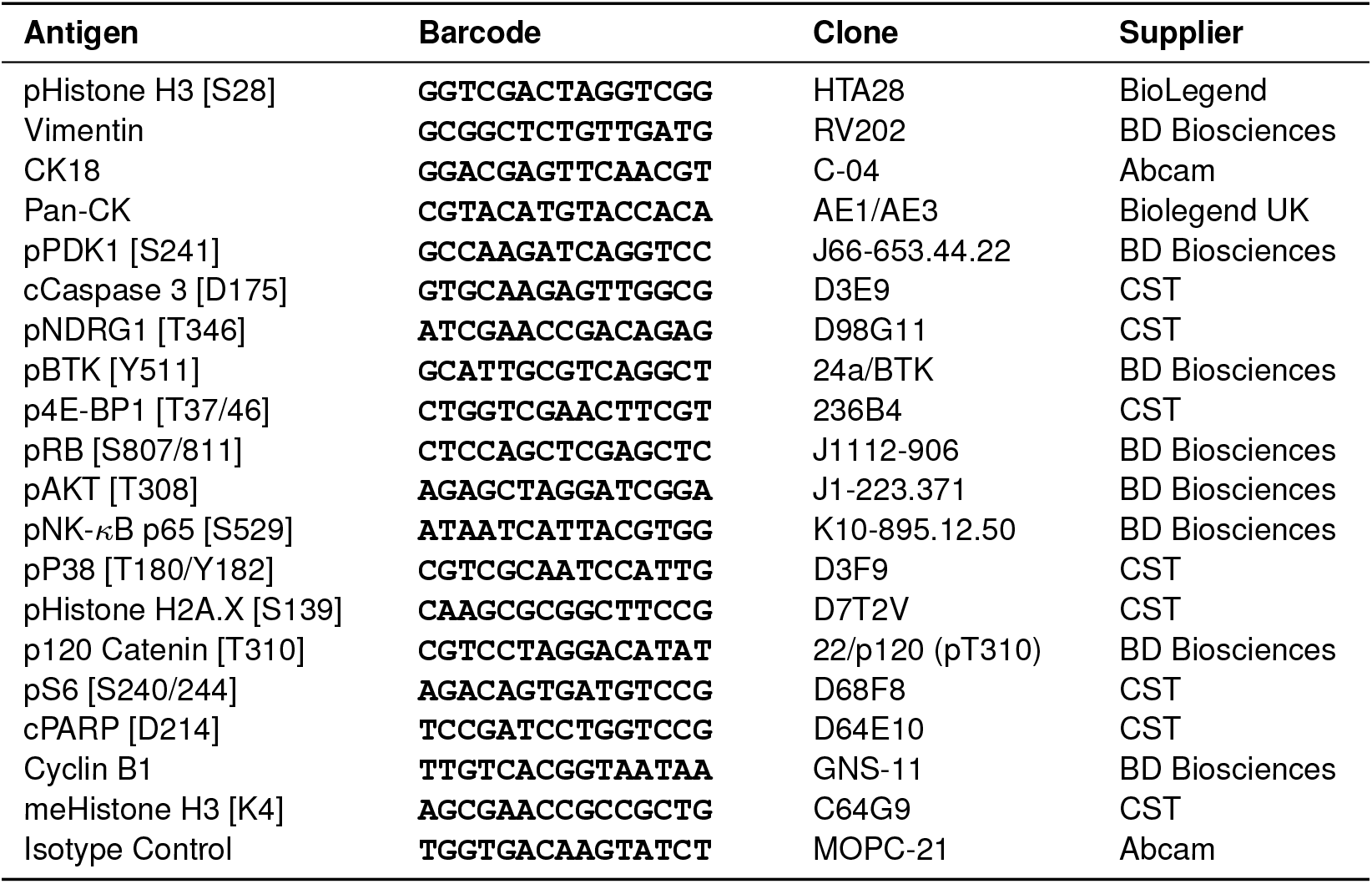
SIGNAL-seq antibody panel used in PDO + CAF organoid experiments.

**Figure 2.**
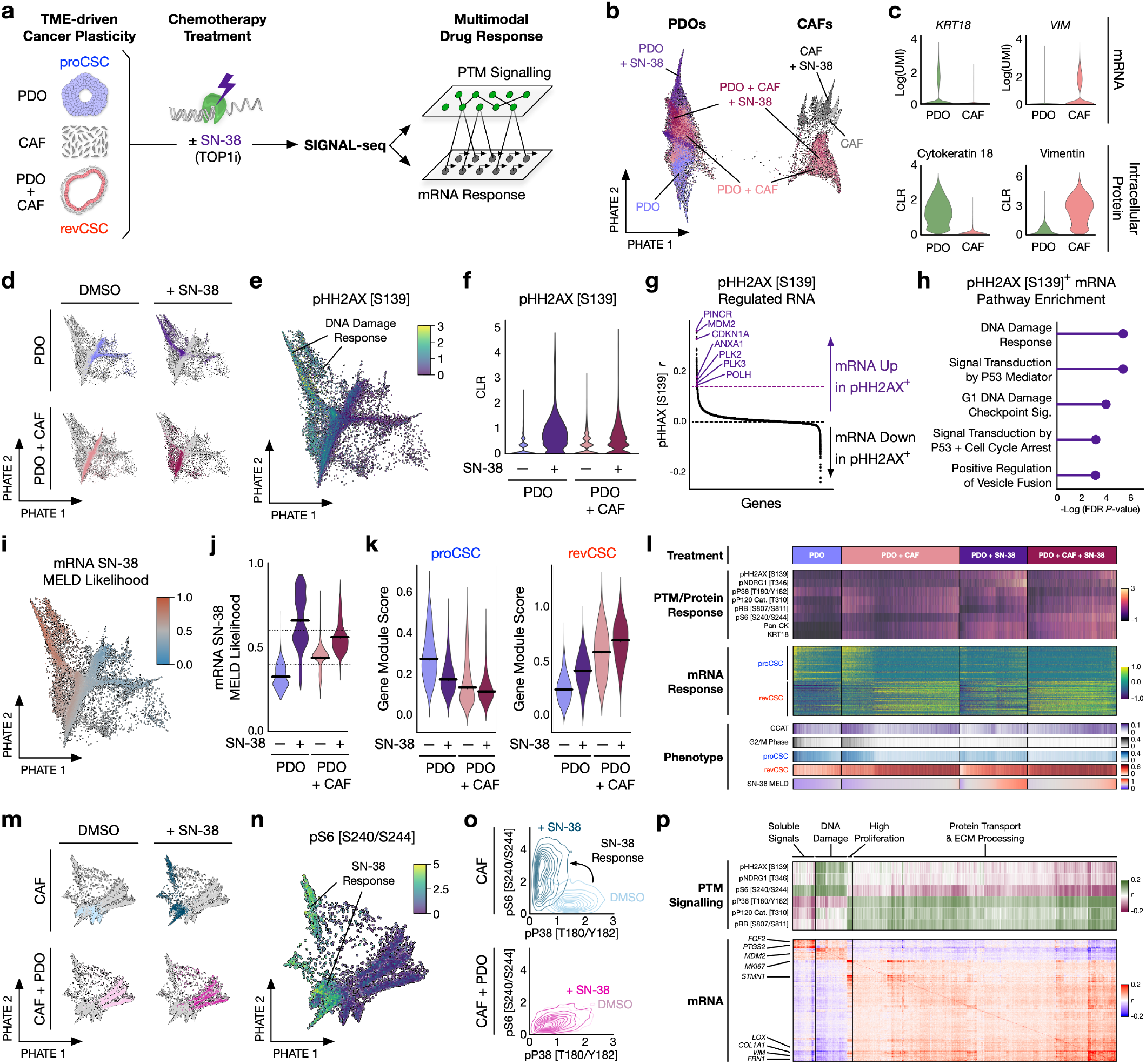
SIGNAL-seq Analysis of Tumour Microenvironment (TME) Organoid Drug Response Identifies Protein PTM Signalling Regulation of Cell Plasticity. **a)**. Experimental workflow. **b)** Single-cell PHATE coloured by experimental condition resolves PDOs and CAFs (30,892 cells) built on RNA modality. **c)** mRNA/protein expression of *KRT18*/Cytokeratin 18 and *VIM*/Vimentin in PDOs and CAFs. **d)** Single-cell PHATEs of PDOs coloured by +/-CAFs, +/-SN-38 built on RNA modality. **e)** PDO cells from all conditions coloured by CLR pHH2AX [S139] detection. **f)** PDO CLR pHH2AX [S139] detection +/-CAFs, +/-SN-38. **g)** PDO RNA gene expression from all conditions correlated against CLR pHH2AX [S139] detection. Black dotted line represents 0, purple dotted line shows threshold for top correlated genes used in **h. h)** RNA gene GO-BP pathway enrichment in the top genes highly correlated with pHH2AX [S139]. **i)** Single-cell RNA PHATE of PDOs coloured by SN-38 MELD likelihood computed using RNA modality. **j)** PDO SN-38 MELD likelihood +/-CAFs, +/-SN-38. Bars represent mean MELD likelihood. **k)** PDO proCSC and revCSC gene signatures +/-CAFs, +/-SN-38. Bars represent median gene module score. **l)** Single-cell heatmap of PDO scaled PTM/protein detection (top - magma color bar), scaled RNA transcripts from pro/rev CSC signatures (middle - viridis color bar), and a collection of phenotypic gene-based module scores (bottom - assorted color bars) cultures +/-CAFs, +/-SN-38, cells ranked within culture by SN-38 MELD likelihood. **m)** Single-cell RNA PHATEs of CAFs coloured by +/-PDOs, +/-SN-38. **n)** CAFs coloured by pS6 [S240/S244]. **o)** Density plots of CAF pS6 [S240/S244] vs pP38 [T180/Y182] +/-PDOs, +/-SN-38. **p)** Pearson correlation heatmap of PTM (rows) - gene (columns) correlations (top, green-purple color bar) and gene-gene (rows and columns) correlations (bottom, red-blue color bar) correlations across CAF single-cells from mono-cultures +/-SN-38 highlighting the close relationship between PTM signalling and the co-regulated RNA response modules.

Following data QC, SIGNAL-seq identified 30,895 single cells and could clearly resolve Cytokeratin 18^+^ PDOs and Vimentin^+^ CAFs at both mRNA and protein levels (Figure 2b, c). Cytokeratin 18 and Vimentin protein measurements are noticeably less sparse than their respective mRNA transcripts — highlighting the value of SIGNAL-seq to measure cell-type-specific intra-cellular proteins.

SIGNAL-seq detected major shifts across both PTM signalling and RNA response in PDOs +/-CAFs, +/-SN-38 (Figure 2d). SN-38 inhibits topoisomerase I, resulting in stalled DNA-replication and DNA-damage [22]. In agreement, SIGNAL-seq detected pHH2AX [S139]^+^ DNA-damage in SN-38 treated PDOs (Figure 2e, f) and identified the transcriptional response of organoid cells actively undergoing DNA-damage (Figure 2g) — including canonical DNA-damage response transcriptional programs that were significantly correlated with pHH2AX [S139] (Figure 2h).

SIGNAL-seq also confirmed that pHH2AX [S139]^+^ proCSC PDOs experience a larger transcriptional response to SN-38 compared to revCSC PDOs (Figure 2i-k). When treated with SN-38 PDO mono-cultures have high levels of pHH2AX [S139], pNDRG1 [T346], and pP38 [T180/Y182], and lose their transcriptional polarisation to proCSC. By contrast PDOs co-cultured with CAFs before SN-38 treatment still experience on-target pHH2AX [S139] DNA-damage, but have low pNDRG1 [T346] and pP38 [T180/Y182], high pS6 [S240/S244] signalling, and are transcriptionally polarised to the chemorefractory revCSC fate (Figure 2l).

CAFs also have distinct PTM and RNA responses to SN-38 and PDOs (Figure 2m). For example, SN-38 switches CAFs from a pP38 [T180/Y182] to pS6 [S240/S244] dominant signalling state in mono-culture, but when cocultured with PDOs CAF signalling can not be perturbed by SN-38 (Figure 2n, o).

By directly comparing PTM and RNA levels with genegene correlation derived modules, SIGNAL-seq revealed that PTM signalling and transcriptional regulation are highly coordinated in CAFs. These PTM and RNA coordinated processes align with key CAF phenotypes such as ECM processing and the secretion of soluble ligands – both of which are known regulators of cancer cell plasticity in the TME [21] (Figure 2p).

By measuring both PTMs and mRNA ligand-receptor pairs in single cells, SIGNAL-seq enables duel intra- and inter-cellular analysis of heterocellular signalling in a single assay. We used SIGNAL-seq data to build a heterocellular signalling model whereby CAF ligands (e.g. FGF7, VEGFC, TGF-β1, HGF) bind to cancer cell receptors (e.g. NRP1, ITGA9, TGF-βR3, MET) to regulate cancer cell PTM signalling (e.g. pP38 [T180/Y182], p120 Catenin [T310]), that drives a transcriptional plasticity response (via down-regulated inferred transcription factor activity of LEF1, ASCL2, and SOX9 and upregulated YAP, AP1, and NF-*κ*B). This transcriptional response polarises chemosensitive proCSC cancer cells to a chemorefractory revCSC fate (Figure 3) (Figure S3). This demonstrates that SIGNAL-seq data can drive an inter- and intra-cellular model describing stromal regulation of cancer cell plasticity in a single assay.

**Figure 3.**
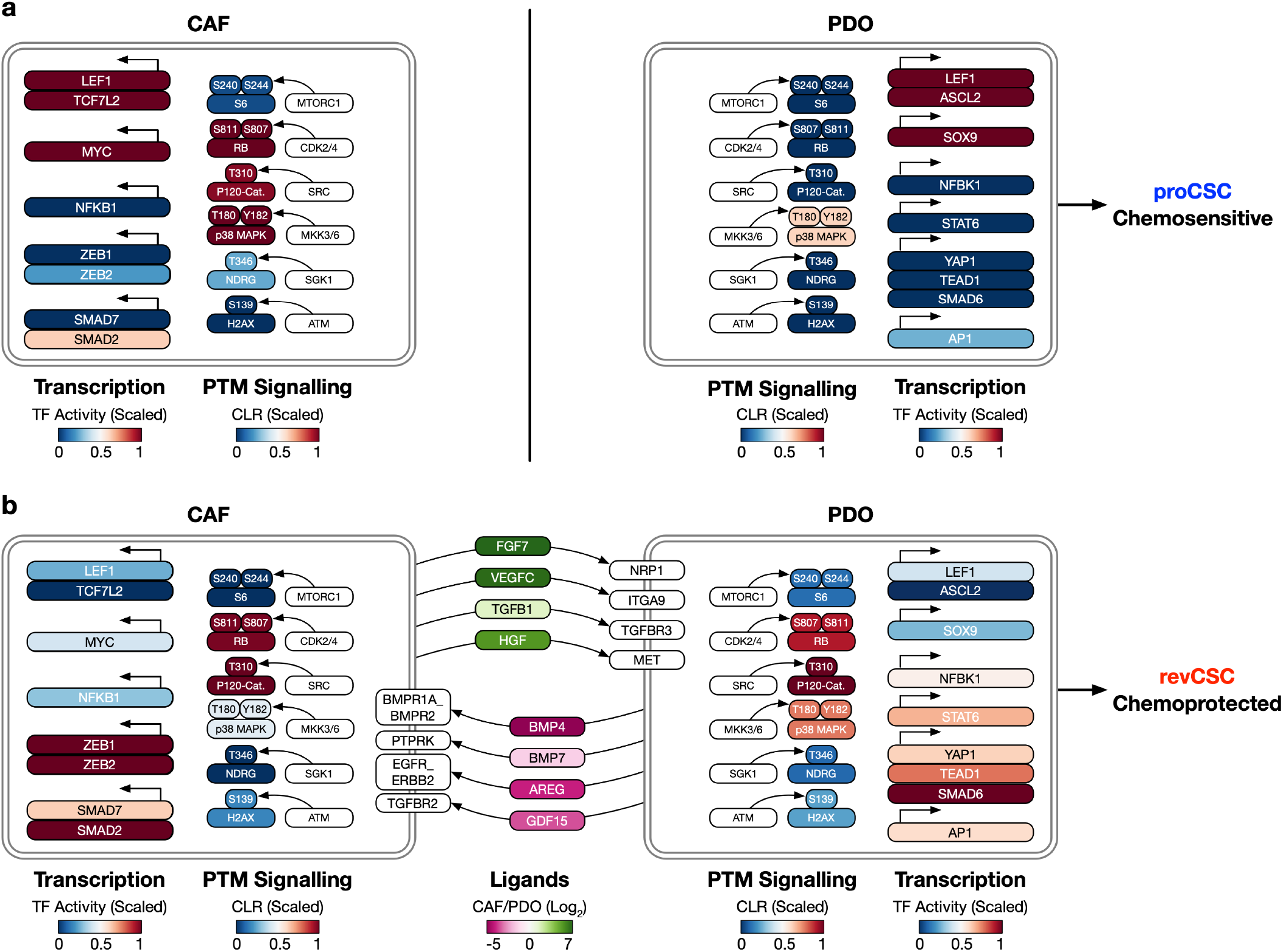
SIGNAL-seq Inter- and Intra-cellular Signalling Analysis of CAF-chemoprotection in CRC PDOs. **a)** Cell-type-specific summary models of PTM signalling (CLR) and inferred transcription factor activity activity of CAF and PDO mono-cultures. **b)** Summary model of inter-cellular communication using mRNA ligand-receptor pairing combined with intra-cellular PTM and inferred transcription factor activity signalling in PDO-CAF co-cultures.

In summary, we demonstrate that combining mass cytometry staining protocols with anti-PTM oligo-tagged antibodies and split-pool combinatorial barcoding enables simultaneous measurement of RNA and intra-cellular PTMs in single cells. SIGNAL-seq does not require specialised or propriety equipment, and is based on a split-pool combinatorial barcoding foundation that scales to hundreds of thousands of cells. Split-pool combinatorial barcoding technology has recently been demonstrated to scale to 1 million cells in a single assay with Perturb-seq CRISPR screening, enabling a step-change in the perturbational space that can be explored [23]. By measuring both mRNA ligand-receptor pairs, PTMs, and transcriptional responses, SIGNAL-seq can assess both inter- and intra-cellular regulation of cell plasticity in a single assay. Given the importance of inter- and intra-cellular signalling across healthy and diseased tissues, we propose SIGNAL-seq as a powerful technology for studying cell-cell communication in heterocellular systems.

## Methods

### Protocols

A step-by-step protocol for SIGNAL-seq is available at tape-lab.com/resources.

Questions regarding the SIGNAL-seq protocol can be submitted to.

### Cell Culture

HeLa cells were obtained from ATCC and cultured in DMEM high-glucose with L-Glutamine and Sodium pyruvate (Thermo Fisher Scientific #41966-029), supplemented with 2 mM L-Glutamine (Sigma #G7513) and 10% FBS (Pan-Biotech #P30-8500). HeLa spheroids were generated by seeding 70k cells per well in a Elplasia® 96-well plate (Corning #4446) as described in Madsen *et al*., [15]. HeLa cells were cultured as spheroids for 48 hours and then starved in serum-free media for 4 hours before treatment. After starvation HeLa spheroids were pre-treated with a combination of inhibitors for 10 minutes before growth factor treatment: 100 nM Trametinib (Cayman #16292) and 500 nM GDC0941 Pictilisib (SelleckChem #S186513) or vehicle (DMSO Sigma #D2650). After inhibitor treatment spheroids were treated with a combination of growth factors or vehicle for 30 minutes before fixation in-situ: 100 nM Human IGF1 (Peprotech #100-11) and 100 nM Human EGF (Peprotech #AF-100-15-1mg). HeLa cells were routinely tested negative for Mycoplasma (Promega PK-CA20-700-20).

### Spheroid SIGNAL-seq Sample Processing

Spheroids in the Elplasia plate were then placed on ice and fixed *in situ* for 20 minutes on ice in 1% effective Formaldehyde added to the serum-free media (Formaldehyde, 16%, methanol-free, Ultra Pure; Polysciences #18814-20). The cell-fixation media was then removed and the cells were incubated with 50 mM TrisHCL (Thermo Scientific #J22638.AE) for 5 minutes on ice and re-suspended in DPBS with Protectorase (Roche #3335402001) and Superase (Invitrogen #AM2694) RNAse inhibitors added at 0.1 U/µL each (RNAse inhibitor cocktail +RI). Spheroids were then re-suspended in 600 µL of PBS +RI and 0.1% UltraPure BSA (Invitrogen #AM2618) and loaded into TissueGrinder tubes (FastForward Technologies) for non-enzymatic, mechanical dissociation [15]. The TissueGrinder protocol “Harsh” was run to dissociate the spheroids. The cells were then pelleted and re-suspended in PBS/BSA plus RNAse inhibitor cocktail. Cells were then permeabilised in 0.2% Triton X-1000 (Sigma #X100) for 3 minutes and re-suspended in 0.5x PBS with 5% DMSO and frozen at -80°C until the SIGNAL-seq barcoding process was performed as described below. Sublibraries with a target of 2200 cells were generated at the lysis stage and 1 was taken forwards for library preparation. The library was amplified with 11 PCR cycles during the second cycling phase of the cDNA amplification step before separating the RNA and ADT libraries by SPRI size selection. The ADT library was then amplified for 10 PCR cycles to build the final Illumina compatible i7 indexed ADT library. We sequenced the RNA and antibody-oligo libraries separately. The ADT library was sequenced on a MiSeq (Illumina) using a MiSeqv3 150bp kit and the RNA library on a NextSeq 550 (Illumina) using a mid-output 150bp kit with the cycle configuration 74 bp Read 1, 6 bp i7, 0 bp i5, 86 bp Read 2 across both platforms.

### Spheroid TOB*is* Mass Cytometry

Spheroids were treated and dissociated into single cells as described above and frozen for storage. An aliquot of frozen cells was thawed and analysed by SIGNAL-seq and the another matched aliquot by thiol-organoid barcoding *in situ* (TOB*is*) mass cytometry (MC) [10, 11]. For the TOB*is* MC analysis, cells were first multiplexed with 3-barcodes (TOB*is*33, TOB*is*34, TOB*is*35) of a 35-plex TOB*is* (7-choose-3) barcode set in suspension using identical concentrations for 2 hours at room temperature before 2x washes with GSH-CSB (1 mM Gulutathione (Sigma #G6529)), Cell Staining buffer (Standard Bio Tools #201068)) and 2x washes with CSB. Cells were then pooled and stained with a panel of rare-earth metal conjugated antibodies (Table 2) comprising identical clones and concentrations to the oligo-conjugated antibodies used in SIGNAL-seq (Table 3). Cells were analysed using a Helios Mass Cytometer (Fluidigm). Data was uploaded to Cytobank and analysed using CyG-NAL (https://github.com/TAPE-Lab/CyGNAL). A detailed step-by-step TOB*is* MC protocol can be found in Sufi *et al*., Nature Protocols, 2021 [11]. CyTOF data was asinh transformed with a cofactor of 5 and preprocessed based on gaussian gating parameters, selecting HeLa cells by gating on Pan-CK^+^EpCAM^+^.

### PDO and CAF Culture

CRC PDO HCM-SANG-0270-C20 was obtained from the Human Cancer Models Initiative (Sanger Institute, Cambridge, UK) and expanded in 12-well plates (Helena Biosciences #92412T) in x3 25 µL droplets of Growth Factor Reduced Matrigel (Corning #354230) per well with 1 mL of Advanced DMEM F/12 (Thermo #12634010) containing 2 mM *L*-glutamine (Thermo #25030081), 1 mM N-acetyl-*L*-cysteine (Sigma #A9165), 10 mM HEPES (Sigma #H3375), 500 nM A83-01 (Generon #04-0014), 10 µM SB202190 (Avantor #CAYM10010399-329 10), and 1X B-27 Supplement (Thermo #17504044), 1X N-2 Supplement (Thermo #17502048), 50 ng mL^−1^ EGF (Thermo #PMG8041), 10 nM Gastrin I (Sigma #SCP0152), 10 mM Nicotinamide (Sigma #N0636), and 1X HyClone Penicillin-Streptomycin Solution (Fisher #SV30010), and conditioned media produced as described in [24] at 5% CO_2_, 37°C. PDOs were dissociated into single cells with 1X TrypLE Express Enzyme (Gibco #12604013) (incubated at 37°C for 20 minutes) and passaged every 10 days. CRC CAFs were a kind gift from Prof. Olivier De Wever (University of Gent). CAFs (ATCC #CRL-1459) were cultured in DMEM (Thermo #11965092) enriched with 10% FBS (Gibco 10082147), and 1X HyClone Penicillin-Streptomycin Solution (Fisher #SV30010) at 5% CO_2_, 37°C. PDOs and CAFs were routinely tested negative for Mycoplasma (Lonza #LT07-701).

### PDO and CAF Co-Culture and Treatment

On Day 0 CRC PDOs were dissociated into single cells as described above, and expanded in 12-well plates in Growth Factor Reduced Matrigel with 1 mL per well of Advanced DMEM F/12, supplemented with 2 mM L-glutamine, 1 mM N-acetyl-l-cysteine, 10 mM HEPES, 1X HyClone Penicillin-Streptomycin Solution, 1X B-27 Supplement, 1X N-2 Supplement, 50 ng mL^−1^ EGF, 10 nM Gastrin I, 10 mM Nicotinamide, 500 nM A83-01 and 10 µM SB202190 at 5% CO_2_, 37°C for 96 hours. On Day 1, CAFs were split into a new flask containing a low-serum media (DMEM supplemented with 2% FBS and 1X HyClone-Penicillin Streptomycin Solution). On Day 4, PDO culture media was changed to a reduced media; Advanced DMEM F/12 supplemented with 2 mM glutamine, 1 mM N-acetyl-L-cysteine, 10 mM HEPES, 1X B-27 Supplement, 1X N-2 Supplement, 10 mM Nicotinamide, 1X HyClone-Penicillin Streptomycin Solution and 25 ng mL^−1^ EGF). PDOs and CAFs were seeded on Day 5 in 96-well plates (Helena Biosciences #92696T) in 50 µL Growth Factor Reduced Matrigel with 300 µL of reduced media. PDOs were seeded at a density of approximately 1.5 × 10^3^ organoids/well and CAFs at 3 × 10^5^ cells/well. PDOs and CAFs were either seeded in mono-culture alone, or in co-culture by mixing together in Matrigel on ice. Cultures were maintained for 72 hours at 5% CO_2_, 37°C, with media changes every 24 hours. On Day 6 and 7, media was replaced with reduced media containing 15 nM SN-38 (Sigma #H0165) or DMSO (Merck #D2650), as a vehicle control. On Day 8, after 72-hours in culture and 48-hours of treatment, cultures were processed for SIGNAL-seq (see below).

### Organoid SIGNAL-seq Sample Processing

PDOs were extracted from Matrigel with ice cold PBS. PDOs were then digested to a single-cell suspension with TrypLE Express Enzyme (Gibco #12604013) for 10 minutes at 37°C on a heated orbital shaker at 300rpm. Cells were then re-suspended in 1 mL PBS + RNAse inhibitor cocktail (RI). Fixation and permeabilisation was then carried out with a modified SPLiT-seq protocol. Single-cell suspensions were fixed with on ice in 1% effective Formaldehyde for 10 minutes before the cells were permeabilized for a further 3 minutes with 0.2% effective Triton X-100. Fixation was then quenched with 50 mM of Tris-HCL (Invitrogen #15568025) before the cells were re-suspended in 300 µL of 0.5x PBS +RI +5% DMSO before freezing and storage at -80°C. After SIGNAL-seq barcoding sublibraries of 9000 cell target input were generated at the lysis stage and 4 were taken forwards for library preparation. The sublibraries were amplified with 8 PCR cycles during the second cycling phase of the cDNA amplification step before separating the RNA and ADT libraries by SPRI size selection. The ADT library was then amplified for 11 PCR cycles to build the final Illumina compatible i7 indexed ADT libraries. We sequenced the RNA and ADT libraries as a pool at a 15:85 ADT:RNA library ratio. i7 sublibray indexes 76-79 were assigned to the RNA modality sublibraries and indexes 80-83 were assigned to the ADT modality sublibraries, matched in numerical order i.e (76-RNA:80-ADT = sublibrary 1). The library pool was sequenced on a NovaSeq (Illumina) using an S2 v1.5 200bp kit with the cycle configuration 85 bp Read 1, 6 bp i7, 0 bp i5, 87 bp Read 2.

### Oligo-Antibody Conjugation

We advise users to select monoclonal antibodies for SIGNAL-seq carefully and validate antibody specificity using defined cues (e.g. purified growth factors or signalling agonists) and specific inhibitors that canonically regulate both protein and PTM antigens when possible. PTM signalling is a dynamic process and we advise users to perform time course analyses in their model systems when establishing anti-PTM antibody panels. For this study, all antibodies were first tested for antigen binding in their respective model systems using TOB*is* mass cytometry [10, 11]. Antibodies were purchased carrier-free and conjugated using the protocol outlined in CITE-seq [3] and subsequently updated and accessed at https://cite-seq.com/protocols, that is based on an original conjugation protocol developed for immuno-PCR [25]. All oligo sequences were ordered from Integrated DNA Technologies (IDT) and had the structure: /5AmMC12/CTACACGACGCTCTTCCGATCT-(contd) XXXXXXXXXXXXXXXBAAAAAAAAAAAAAAAAAA*A*A where the construct is designed beginning with a partial Illumina TruSeq read 1 primer that is used as the PCR handle and the X is representative of the antibody specific 15 nt barcode. Complete antibody conjugation was validated by running both conjugated and un-conjugated paired antibodies on a 4-15% SDS-PAGE gel (Bio-Rad #4561083) under non-reducing conditions. Confirmation of the antibody cleanup steps to remove free (un-conjugated) ssDNA oligo was performed using a 4% agarose gel. All antibody concentrations were determined using a BCA assay (Thermo #23227). Antibodies were pooled individually after antibody free oligo cleanup for the experiment in Figure 1 and then pooled together before cleanup (Table 3) or pooled individually after cleanup for the experiment in Figure 2 (Table 4).

### SIGNAL-seq Oligo-Antibody Staining

Cells were thawed at 37°C until a small ice crystal remained and then incubated on ice. Cells were then added to a 96-well Nunc M Well Plate (Thermo #267334) and washed with blocking buffer; PBS + 1% BSA, 0.1% Tween (Thermo #85113), 0.025% Dextran Sulfate (Thermo #D8906), 1 mg mL^−1^ Salmon Sperm DNA (Thermo #15632011), 1:100 FcX TruStain (Biolegend #422301) with RNAse inhibitor (RI) cocktail. Cells were then incubated in blocking buffer on ice for 15 minutes after which the cells were stained in a total volume of 75 µL with Oligo-antibody cocktail in blocking buffer + RI for 30 minutes at room temperature with mixing by gentle pippetting every 10 minutes with a wide-bore tip. The antibody panel for the spheroid experiment in Figure 1 is described in Table 3 and the antibody panel for PDO+CAF experiment in Figure 2 and Figure 3 is described in Table 4.

Cells were then washed 2x with blocking buffer and 1x with SIGNAL-seq wash buffer (PBS +0.5%BSA, 0.1% Tween + RI) before re-suspension in 0.5x PBS + RI. Cells were then filtered two or more times through a 40 µm FlowMi filter (Flowmi #BAH136800040) until no cell multiplets could be seen under a microscope and counted. Cells were then re-suspended to the appropriate concentration in 0.5x PBS + RI for loading into the SIGNAL-seq reverse transcription barcoding plate.

### SIGNAL-seq Split-pool Barcoding

Split-pool barcoding combinatorial indexing of both transcriptome and antibody-oligo tags (ADT) were performed as per the SPLiT-seq protocol [9] with minor modifications. An updated barcode plate setup described in the Micro-SPLiT method [26] was used. A complete list of oligos is provided in the online Supplementary Material. All oligo sequences were ordered from Integrated DNA Technologies (IDT) other than the Template Switching Oligo (TSO -BC_0127) which was purchased from Qiagen. Briefly, after staining with Oligo-Antibodies, cells were diluted in 0.5x PBS + RI and 8 µL loaded into 12 µL of RT-mix in the 48-wells of the SPLiT-seq barcode-1 reverse transcription plate and reverse transcription performed with Maxima H Minus Reverse Transcriptase (Fisher Scientific #EP0753). Cells were then pooled and resuspended into NEB 3.1 Buffer (New England Bio-Labs #B7203S), T4 DNA Ligase (New England BioLabs #M0202L) and Ligase Buffer (New England BioLabs #B0202S). Cell-ligase reaction mixture (40 µL) was then loaded into the 96-well barcode-2 ligation plate (10 µL) and ligation was carried out in a thermocycler for 30 min at 37°C. The round 2 blocking solution (10 µL) was added to each well of the barcode-2 plate and incubated in a thermocycler for 30 min at 37°C. Cells were then pooled and filtered through a 40 µM filter into a basin and 100 µL T4 DNA ligase was added. 50 µL Cell-ligase reaction mixture was then loaded into the 96-well barcode-3 ligation plate (10 µL) and ligation was carried out in a thermocycler for 30 min at 37°C. The round 3 blocking solution (20 µL) was added to each well of the barcode-3 plate and cells were then pooled into a new basin and filtered through a 40 µM into a 15 mL tube on ice. Cells were then washed with 4 mL 0.1% Triton X-100 solution and resuspended in 50 µL of 1x PBS +RI as described in [9]. All temperature incubation steps were performed in a thermocycler to increase temperature stability. At the end of the barcoding process cells were counted on a haemocytometer using DiYo (Stratech #17579) with a GFP filter on an EVOS FL microscope to resolve in-tact cells from debris. Cells were diluted in 1x PBS +RI and split into 25 µL sublibraries, combined with 25 µL of 2x SPLiT-seq lysis buffer with 5 µL of Proteinase K (Thermo #AM2546) per sublibrary, then incubated at 65°C for 60 minutes and stored at -80°C until library preparation and sequencing. Complete lysis of cells was visually verified by observing DiYo-stained lysates under an EVOS FL microscope using a GFP filter.

### SIGNAL-seq Library Preparation and Sequencing

We performed cDNA isolation and amplification based on the SPLiT-seq protocol [9] with some modifications, a full protocol is provided in the Protocols section above. We performed all steps in the SPLiT-seq library preparation protocol scaled down to half volume to enable thermocycler compatibility and reduce reagents quantity used, with the exception that each sublibrary was incubated with a total effective volumetric quantity of 44 µL of myOne C1 Dynabeads (Thermo #65001), re-suspended in 55 µL of 2xBW+RI and processed at as described in Roseneberg *et al*. [9]. We selected the cDNA amplification cycle numbers based on initial cycle number optimisation of a range of cycles using EvaGreen dye qPCR Ct saturation and pilot sequencing of smaller 250-350 cell sublibraries on an Illumina MiSeq platform to inform cycle selection of a particular sublibrary cell quantity and sample type combination. In order to selectively amplify the antibody-oligo derived tag library we spiked in 0.9 µL of 2 µM primer specific to the PCR handle region of the antibody oligo tag into the cDNA amplification PCR reaction mix (100 µL reaction volume), with the sequence: (ADT_cDNA) CTACACGACGCTCTTC-CGATCT. After cDNA amplification, a 0.6x SPRI bead (Roche #KK8000) cleanup was performed keeping the supernatant which contains the antibody-oligo library, whilst the RNA library was bound the the beads fraction, which was eluted and processed as described in the SPLiT-seq protocol using the Nextera XT kit (Illumina #FC-131-1024) to generate the final RNA derived cDNA SPLiT-seq libraries [9], we performed a final 0.6-0.8x double sided SPRI library cleanup on the RNA library before quantification, pooling and sequencing. We took forward the antibody-oligo library containing SPRI cleanup supernatant and performed a further 2x rounds of a 2x SPRI cleanup to remove residual PCR primers. We took half of the antibody-oligo quantity forwards to build Illumina compatible libraries. We performed PCR amplification using a standard KAPA HIFI mastermix protocol (Roche #KK2601) using a p5 custom library oligo, with the sequence: (ADT_lib) AATGATACGGCGACCACCGA-GATCTACACTCTTTCCCTACACGACGCTC and one of the SPLiT-seq sublibrary i7 index oligos BC0076-BC0083 combining 45 µL ADT cDNA with 50 µL 2x KAPA HiFi HotStart ReadyMix (Roche #KK2601) and 2.5 µL of 10 µM each PCR primer and ran the ADT-lib PCR cycle outlined in Table 5. We selected the cDNA amplification cycles (with a range of 6-12x total) based on optimisation of cycle number using a spare sublibrary in a similar manner to cDNA amplification step as this step is influenced by the number of cells, antibodies, the sample type and treatment. We purified the the resulting library with a 1.2x SPRI cleanup. Final sequencing libraries were quantified using a Qbit DNA HS kit (Thermo #Q32856) for concentration and a Bioanalayzer HS DNA chip (Agilent #5067-4626) for amplicon size. This ultimately results in an Illumina compatible library structure where the RNA cDNA is present in Read 1, the ADT barcode is also present in Read 1 (bp 1-15). Both the ADT and the RNA libraries have the UMI and barcode sequence in Read 2 with the following structure UMI: 1-10 bp, BC3 (L3): 11-18 bp, BC2 (L2): 49-56 bp, BC1 (RT1): 79-86 bp.

**Table 5.** ADT PCR Cycle. Run time = 25-45 minutes, lid temperature = 105°C, sublibrary volume = 100 µL.

| Step | Time | Temperature |
| --- | --- | --- |
| 1 | 3 Minutes | 95°C |
| 2 | 20 Seconds | 95°C |
| 3 | 30 Seconds | 60°C |
| 4 | 20 Seconds | 72°C |
| *Go to Step 2, repeat 5-11 times (6-12x cycles total)* |  |  |
| 5 | 5 Minutes | 72°C |
| 6 | ∞ | 4°C |

### SIGNAL-seq data Preprocessing and Analysis

Sequencing data was demultiplexed using the sublibrary i7 indexes. FASTQ RNA data was aligned to the GRCh38 reference genome using STAR (v2.7.3a) [27] and processed with samtools (v1.9) [28]. Cell barcodes (CB) reads were collated, Oligo-dT (PolyA) and random Hexamer (rHex) barcodes were merged and UMIs counted using the zUMIs package (v2.9.7) to generate a cell x gene digital gene expression matrix (DGE) for each sublibrary (https://github.com/sdparekh/zUMIs) [29]. CBs were merged based on a total of 2 hamming distance mismatches across the full CB and UMIs were collapsed based on a 1 hamming distance mismatch. We used both intronic and exonic UMI counts. CB barcode well location and sample identity was annotated and initial QC performed with the splitRtools R package (https://github.com/TAPE-Lab/splitRtools). Antibodyoligo data was aligned to an antibody-barcode index using the Kallisto KITE pipeline from the BUStools package (https://github.com/pachterlab/kite) [30] and RT CBs Oligo-dT and rHex barcodes were merged before the ADT barcodes were mapped to their respective RNA sublibrary specific shared CB using the Anndata (v0.10.4) data structure [31], Scanpy (v1.9.6) [32] and Muon (v0.1.5) [33].

### HeLa SIGNAL-seq Data QC and Analysis

RNA modality CBs were first filtered based on the following parameters: CBs >2xIQR of the median log10(UMIs), CBs <500 UMIs, CBs > 12000 genes, CBs <100 genes, CBs >30% mitochondrial UMIs, CBs >0.75 UMI/read ratio and genes that were detected in less than 3 cells. The combined multimodal CB matrix was then further filtered in the ADT modality based on the following parameters: CBs <40 UMIs, CBs <11 antigens detected. The matrix was further filtered based on outliers in isotype control ADT UMIs. The RNA CBs were then matched to ADT CBs and CBs not detected in both modalities were discarded. This ultimately resulted in a multimodal dataset of 1143 cells. RNA modality UMI counts were then normalised in Scanpy with count based normalisation using a scaling factor of 10,000 excluding highly expressed genes with a max_fraction=0.05. The normalized matrix was then natural log transformed for downstream analysis. The ADT modality was centered Log ratio transformed (CLR) using the Muon clr() function. We used the PHATE algorithm [34] with default settings for visualisation purposes of the data using the ADT modality data with a subset of 15 signalling-associated markers as input. Long non-coding RNA identity was determined using LNCipedia gene annotation conversion table v5.2 [35]. EMDs were calculated using the emd() function in scprep (https://scprep.readthedocs.io/en/stable/index.html). pAKT[T308] and pPDK1 [S241] were removed from EMD correlation analysis and PHATE embeddings as their staining profiles in mass cytometry do not match prior knowledge of HeLa signalling dynamics [15]. Early EGF response genes were taken from *Amit et al*. [17] and gene module scoring was performed with the score_-genes function in Scanpy using default settings on Log-Normalised data. PTM-Gene correlations were computed using the Pearson correlation between CLR PTM and Log-Normalised RNA data. RNA modality cell cycle scoring was used based on a list of S-phase and G2M-phase genes previously outlined in *Tirosh et al*. [19] with the Scanpy function score_genes_cell_cycle. For RNA modality benchmarking the SPLiT-seq cell line mixing FASTQ dataset from *Rosenberg et al*. [9] was downloaded from SRA (SRR6750057) and processed as described above for the SIGNAL-seq data, mapped to a combined species reference genome of GRCh38 and mm10. Both datasets were downsampled to 65k reads per cell using the downsampling function in the zUMIs package. HeLa cells were identified from the *Rosenberg et al*. data by selecting the top 3000 highly variable genes (HVG) using the ‘seurat v3 method’ in Scanpy, scaling the data, performing PCA on the HVGs and using the first 30 PCs to build a neighborhood graph, where the cells were clustered using Leiden clustering with default settings and res=0.3 to generate 3 clusters that identify the respective cell lines to subset the HeLa cell line data. Genes per cell were compared between the SPLiT-seq HeLa cells and the cC3^Low^ SIGNAL-seq HeLa cells.

### Organoid SIGNAL-seq data Preprocessing and Analysis

RNA modality CBs were first filtered based on the following parameters: Neotypic doublets between PDO and CAF were identified with Scrublet [36] given an estimated doublet rate of 3% based on previous published data. CBs >9250 UMIs, CBs <500 UMIs, CBs >4000 genes, CBs <300 genes, CBs >15% mitochondrial UMIs, CBs >0.4 UMI/read ratio and genes that were detected in fewer than 25 cells. The combined multimodal CB matrix was then further filtered in the ADT modality based on the following parameters: CBs >5000 UMIs, CBs <75 UMIs, CBs > 18 antigens detected, CBs < 4 antigens detected. The matrix was further filtered based on outliers in isotype control ADT UMIs. CBs were then discarded that did not have both RNA and ADT modality data present. This ultimately resulted in a multimodal dataset of 30,895 cells. RNA modality UMI counts were then normalised in Scanpy using count based normalization with a scaling factor of 10,000 and the normalised matrix was then natural log transformed for downstream analysis. The ADT modality matrix was Centered Log Ratio (CLR) transformed for downstream analysis. To identify epithelial and Fibroblast cell types PCA was performed will all genes as input and a neighborhood graph was constructed with the first 100 PCs as input. Clustering was performed with Leiden clustering on the neighborhood graph with res=0.1 to generate clusters that represented both epithelial PDO and fibroblast. PHATE embeddings of the RNA modality were used for dataset visualization of PDO and CAF co-culture data, regressing out total UMI counts per cell and scaling the data for input into PHATE. For the PHATE embeddings 100 PCs were used with default parameters with modifications for the PHATE of all cells (k=15, t=7), the PHATE embedding of PDO (t=12) and the PHATE embedding of Fibroblasts (t=12). Gene ontology (GO) analysis was performed with the GSEAPY [37] package enrichr [38] module using the GO_Biological_Process_2023 gene sets [39]. MELD [40] was used to identify PDO cell populations within each culture condition that are perturbed based on treatment with SN-38. MELD is a method that uses manifold-geometry to quantify the effect of an experimental perturbation (SN-38 treatment) by estimating the relative likelihood of observing cells in each experimental condition over a graph learned from all cells. We used SN-38 treatment as the perturbation across all PDO using Log-normalised RNA PDO single-cell data as input. Multimodal heatmaps displaying both RNA and ADT data were made with the ComplexHeatmap package [41]. CCAT score [42] was used to calculate signalling entropy and estimate the differentiation potency of our single-cell PDO epithelial data. Highly differentially expressed genes (DEGs) were calculated between Fibroblasts and PDO epithelial cells from the stated conditions using the wilcoxon rank sum test with the scanpy rank_genes_groups function. The gene-gene and PTM-gene correlations were plotted for the top DEGs between CAF mono-cultures selected with a BH adjusted p-value <0.01 and an absolute value Log Fold Change >0.5.

### Generation of proCSC and revCSC Signature Scores

single-cell PDO specific proCSC and revCSC signatures were computed by scoring each PDO epithelial cell for literature derived proCSC and revCSC CRC genes identified by Ramos Zapatero *et al* [12] with the score_genes function in Scanpy using default settings on Log-Normalised data. To derive refined single-cell proCSC and revCSC gene signatures, these two literature proCSC and revCSC gene scores were then correlated against the single-cell gene expression matrix for the control treated PDO epithelial cells to identify a more comprehensive list of genes significantly associated with either CRC stem cell phenotype (r >0.1, >3 SD above the mean for shuffled data). Full gene signatures are detailed in online Supplementary Material. The top 35 correlated genes from each signature were used for visualisation in the heatmap.

### Ligand-Receptor Analysis

Predicted ligand-receptor (LR) interactions between cell types were computed using the liana-py implementation of the LIANA+ framework [43] using the Omnipath ligand-receptor database [44] to generate a candidate list of LR interactions across the dataset. The Liana rank_aggregate function with a minimum LR expressing fraction of the cell population of 0.1 (expr_prop) was used to generate a consensus integrating the predictions of an ensemble of methods; CellPhoneDB [45], Connectome [46], log2FC [43], NATMI [47], SingleCellSignalR [48] and CellChat [49]. The resulting LR interactions were then filtered for those interactions where the parent_intercell_source class of molecule was defined as a ‘Ligand’ to select for ligands which might originate from either cell type. Ligands were then ranked by LogFC on significant differentially expressed ligands using a wilcoxon rank sum test (BH corrected pval <0.05) between PDO epithelial cells in mono-culture and co-culture to identify possible PDO ligands enriched in PDO mono-cuture that might represent autocrine LR interactions contributing to the proCSC PDO phenotype and ligands that were specifically enriched in Fibroblasts in PDO-CAF co-cultures that might contribute to the revCSC PDO phenotype through heterocellular LR signalling.

### Transcription Factor Activity Analysis

Transcription factor downstream pathway activity inference was computed with the decoupler python package [50] using the Univariate Linear Model (ULM) method with the Omnipath CollecTRI knowledge graph network [44], comprising a comprehensive resource of curated transcription factors (TFs) and their transcriptional targets. Briefly, a linear model was fitted that predicts the observed gene expression based on a given TF’s TF-Gene interaction weights, where the t-value of the slope of the fitted model is taken as the TF activity score. TF activities were selected for visualization that had significantly different inferred activity (p_adj<0.05) between PDO, CAF in both mono-culture and co-culture respectively using the decoupler rank_sources_groups function implementing the ‘t-test_overestim_var’ statistical method for computing differences between groups.

### Immunofluorescence

HeLa spheroids were cultured and treated as previously described. All steps were performed with spheroids in Elplasia® 96-well plates. Spheroids were washed with PBS and fixed with 4% ice cold formaldehyde on ice for 1 hour. The spheroids were then permeabilised with spheroid permeabilisation buffer (SPB; 0.2% BSA, 0.1% Triton-X100, PBS) at room temperature for 30 minutes. Spheroids were then blocked with CSB (Standard Bio Tools #201068) and stained with identical antibody clones to the SIGNAL-seq experiment pS6 [S240/S244]-AF488 (CST #5018) 1:100, Cleaved-Caspase-3 [D175]-AF647 (CST #9669) 1:100 overnight at 4°C. Following staining, spheroids were washed 3x with CSB, washed with B/PBS (1% BSA, PBS) and stained with Hoechst (Sigma #14533-100mg) 5 mg/mL stock diluted 1:1000 in B/PBS for 45 minutes at room temperature. Spheroids were then washed 3x times with B/PBS before a final PBS wash before imaging. Spheroids were imaged on an Opera Phenix Plus High-Content Screening System (PerkinElmer, #HH14001000). Images were analysed in Fiji [51].

## Supporting information

SIGNAL-seq Oligos

SIGNAL-seq Gene Signatures

## Data Availability

Raw scRNA-seq data has been deposited at NCBI GEO under the accession GSE256405. Processed RNA and ADT scRNA-seq data and metadata can be accessed at Zenodo: https://doi.org/10.5281/zenodo. 10676523. Rosenberg *et al*. SPLiT-seq data is available for download from SRA under the accession SRR6750057. TOB*is* MC data are available as a Community Cytobank https://community.cytobank.org/cytobank/experiments/114601.

## Code Availability

Code to process SIGNAL-seq data and generate the figures from this paper is available at https://github.com/TAPE-Lab/SIGNAL-seq.

Questions regarding the SIGNAL-seq protocol can be submitted to.

## Acknowledgements

We are extremely grateful to M. Garnett, H. Francies and the Cell Model Network UK for sharing CRC PDOs and O. De Wever for providing CRC CAFs. We thank B. Vanhaesebroeck for access to the TissueGrinder. We thank Y. Guo and the UCL CI Flow Cytometry Translational Technology Platform for mass cytometry support, A. McLatchie and the UCL CI Genomics Translational Technology Platform, and UCL Genomics for sequencing support, and P. Vlckova and the UCL Organoid Translational Technology Platform for organoid culture support. We thank members of Tape Lab and E. Sahai for for their constructive critique of the manuscript.

This work was funded by the UKRI Medical Research Council (MR/T028270/1), Cancer Research UK (C60693/A23783), the Cancer Research UK City of London Centre (C7893/A26233), The Rosetrees Trust (PGS23/100164), and the UCLH Biomedical Research Centre (BRC422).

## Author Contributions

J.O. developed SIGNAL-seq, performed all experiments, analysed the data, and wrote the paper. R.O.S. performed organoid experiments. J.S. developed oligo- and metal-conjugated antibody panels. R.M. established the HeLa spheroid assay. X.Q. provided sequencing support. E.B. provided organoid support. C.J.T. conceived SIGNAL-seq, designed the study, analysed the data, and wrote the paper.

## Supplementary Information

### Supplementary Figures

**Figure S1.**
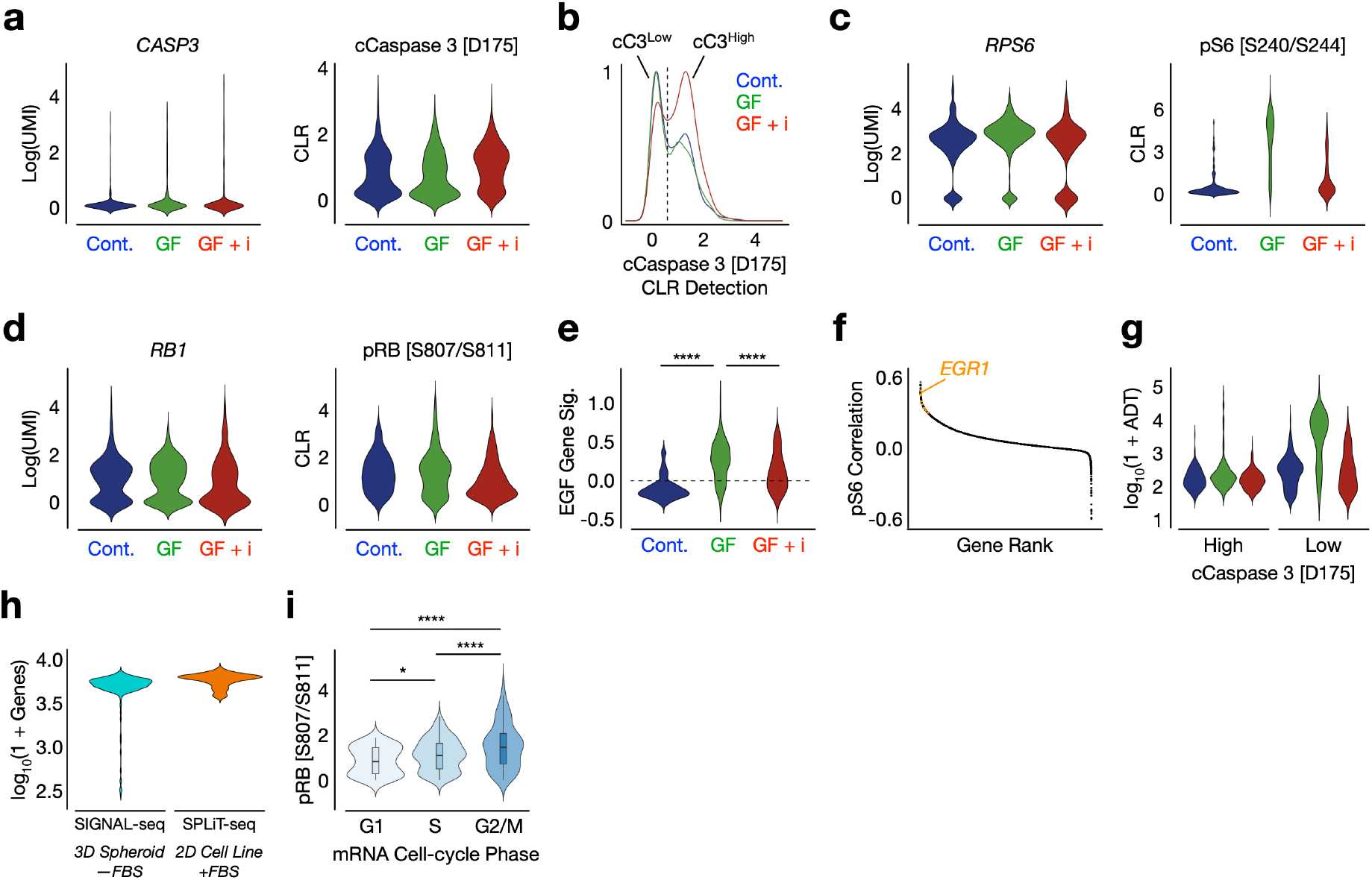
SIGNAL-seq Analysis of Perturbed 3D Spheroids. **a)** SIGNAL-seq detection of *CASP3* mRNA and cCaspase 3 [D175] in control, +IGF +EGF (+GF), or +IGF +EGF +MEKi + PI3Ki (GF+i) 3D spheroids. **b)** Identification of viable (cCaspase 3 [D175]^Low^) and apoptotic (cCaspase 3 [D175]^High^) cells. **c)** *RPS6* mRNA and pS6 [S240/S244] in cCaspase 3 [D175]^Low^ cells. **d)** *RB1* mRNA and pRB [S807/S811] in cCaspase 3 [D175]^Low^ cells. **e)** EGF HeLa mRNA response gene signature module score from [17]. **f)** Gene rank vs Pearson correlation plot of gene -pS6 [S240/S244] correlation, genes from the EGF gene signature are highlighted in orange. **g)** Antibody derived tags (ADTs) UMIs per cell identified across each condition, in cCaspase 3 [D175]^High^ and cCaspase 3 [D175]^Low^ cells. **h)** Genes detected per cell after downsampling both datasets to 65k reads per cell in SIGNAL-seq analysis of 3D HeLa spheroids and SPLiT-seq analysis of 2D HeLa cell line grown in serum [9]. **i)** pRB [S807/S811] detection per cell grouped by mRNA predicted cell-cycle phases. t-test, using Holm method for multiple comparison testing = * <0.05, **** <0.0001.

**Figure S2.**
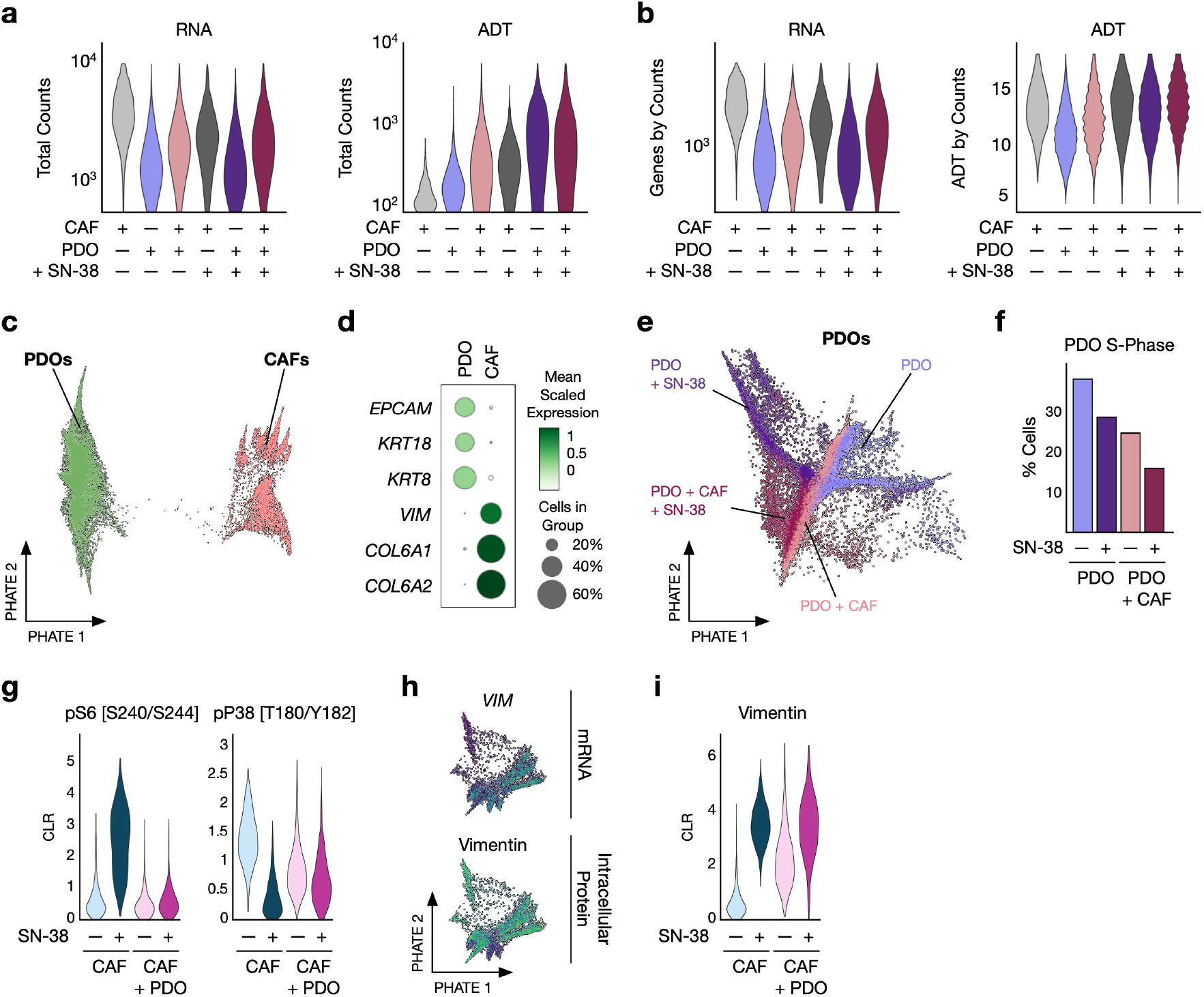
SIGNAL-seq Analysis of Patient-Derived Organoids and Cancer Associated Fibroblasts During Therapy. **a)** Total RNA and ADT counts for PDOs and CAFs +/-SN-38. **b)** Total genes and ADTs per count for PDOs and CAFs +/-SN-38. **c)** Single-cell PHATE of all cells coloured by PDO and CAF clusters. **d)** Canonical epithelial and mesenchymal genes expression per cell-type cluster. **e)** Single-cell PHATE of all PDO cells coloured by treatment. **f)** mRNA estimate of the proportion of PDO cells in S-phase +/-CAFs +/-SN-38. **g)** CAF pS6 [S240/S244] and pP38 [T180/Y182] +/-PDOs +/-SN-38. **h)** Single-cell PHATE of all CAFs coloured by *VIM* mRNA and Vimentin protein. **i)** Vimentin protein levels in CAFs +/-PDOs +/-SN-38.

**Figure S3.**
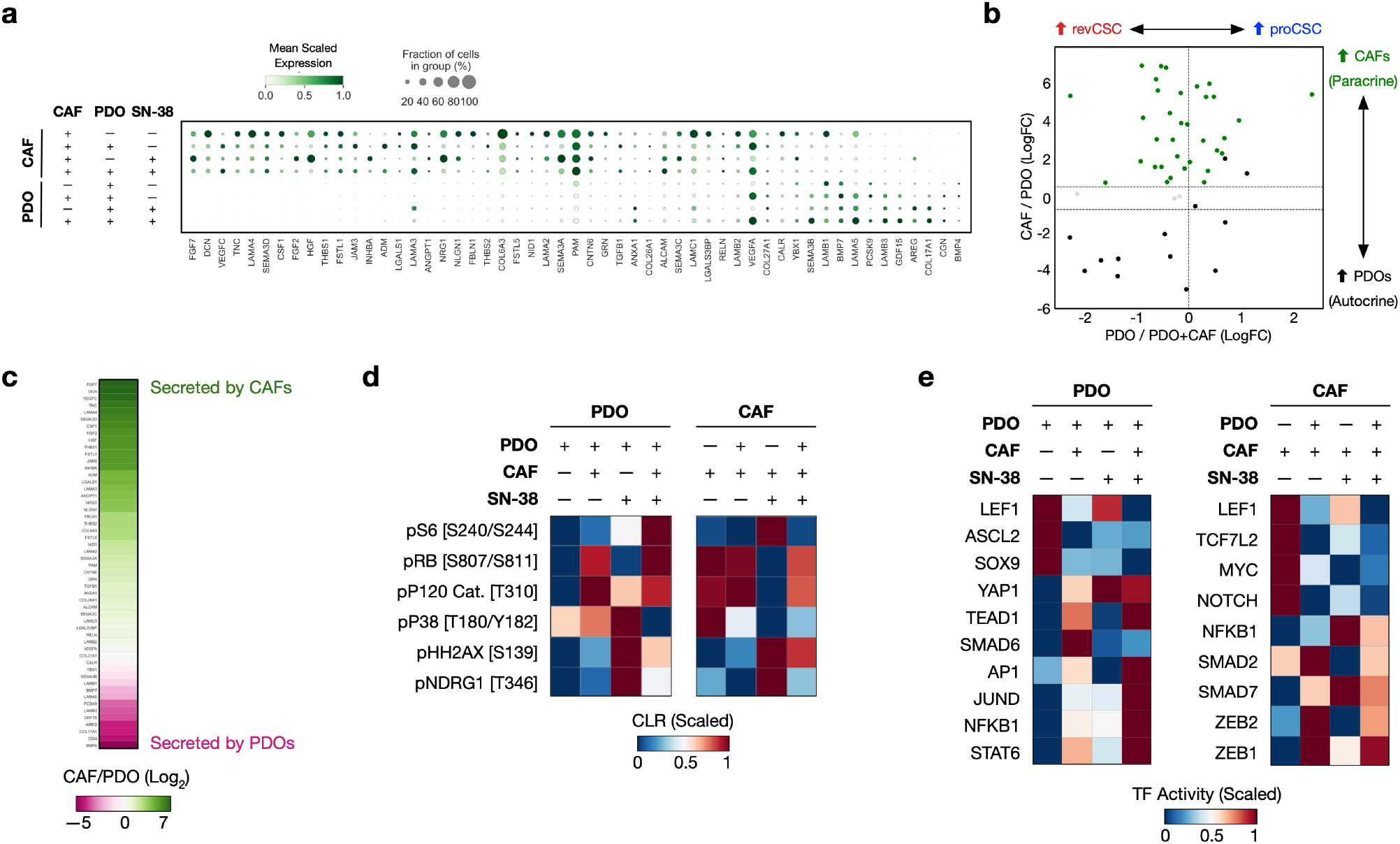
SIGNAL-seq Inter- and Intra-cellular Signalling Analysis. **a)** Ligands expressed by CAFs and PDOs +/-co-culture, +/-SN-38. **b)** CAF/PDO vs PDO/ PDO +CAF ligand expression. **c)** CAF/PDO ligand expression. **d)** PTM signals in PDOs and CAF +/-co-culture, +/-SN-38. **e)** Transcription factor activity in PDOs and CAF +/-co-culture, +/-SN-38.

